# TPLATE complex dependent endocytosis is required for shoot apical meristem maintenance by attenuating CLAVATA1 signaling

**DOI:** 10.1101/2022.10.16.511936

**Authors:** Jie Wang, Qihang Jiang, Roman Pleskot, Peter Grones, Grégoire Denay, Carlos Galván-Ampudia, Elmehdi Bahafid, Xiangyu Xu, Michael Vandorpe, Evelien Mylle, Ive De Smet, Teva Vernoux, Rüdiger Simon, Moritz K. Nowack, Daniel Van Damme

## Abstract

Endocytosis regulates the turnover of cell surface localized receptors, which are crucial for plants to sense and rapidly respond to both endogenous and environmental stimuli. The evolutionarily ancient TPLATE complex (TPC) plays an essential role in clathrin-mediated endocytosis (CME) in Arabidopsis plants. Knockout or strong knockdown of single TPC subunits causes male sterility and seedling lethality phenotypes, complicating analysis of the roles of TPC during plant development. Partially functional alleles of TPC subunits however only cause very mild developmental deviations. Here, we took advantage of the recently reported partially functional TPLATE allele, WDXM2, to investigate a role for TPC-dependent endocytosis in receptor-mediated signalling. We discovered that reduced TPC-dependent endocytosis confers a hypersensitivity to very low doses of CLAVATA3 (CLV3) peptide signalling. This hypersensitivity correlated with the abundance of the CLV3 receptor protein kinase CLAVATA1 (CLV1) at the plasma membrane. Genetic analysis and live-cell imaging revealed that TPC-dependent regulation of CLV3-dependent internalization of CLV1 from the plasma membrane is required for CLV3 function in the shoot. Our findings provide evidence that clathrin-mediated endocytosis of CLV1 is a mechanism to dampen CLV3-mediated signaling during plant development.

## Introduction

Coordinating cellular responses to environmental stimuli largely relies on receptor-like kinases (RLKs) or receptor-like proteins (RLPs) localized on the plasma membrane (PM), that are activated by cognate peptide ligands (Claus *et al*, 2018; Gou & Li, 2020; Hohmann *et al*, 2017; Olsson *et al*, 2019). CLAVATA1(CLV1)-type receptors are one of the most intensively studied groups of plant RLKs, and they are crucial for shoot apical meristem (SAM) and root apical meristem (RAM) maintenance (Clark *et al*, 1993; Clark *et al*, 1997; DeYoung *et al*, 2006; Deyoung & Clark, 2008; Dievart *et al*, 2003; Stahl *et al*, 2013). PM abundance and vacuolar targeting of CLV1 depends on the CLAVATA3 (CLV3) peptide (Nimchuk *et al*, 2011). However, how CLV1 signaling is modulated by its internalization remains unknown (Yamaguchi *et al*, 2016).

In plants, clathrin-mediated endocytosis (CME) is the best-characterized pathway by which cells internalize transporters, receptors and their bound ligands from PM via transport vesicles (Paez Valencia *et al*, 2016; Zhang *et al*, 2015). Internalization of PM localized receptors can occur in a ligand-independent or ligand-dependent manner (Beck *et al*, 2012; Ben Khaled *et al*, 2015; Irani *et al*, 2012; Mbengue M, 2016; Nimchuk *et al*., 2011; Ortiz-Morea *et al*, 2016) and serves either to attenuate signalling by vacuolar degradation or to sustain signalling from endosomes (Claus *et al*., 2018; Paez Valencia *et al*., 2016).

The heterotetrameric adaptor protein complex 2 (AP-2) and the octameric TPLATE complex (TPC) jointly function as adaptor complexes to execute CME in plants (Di Rubbo *et al*, 2013; Gadeyne *et al*, 2014; Zhang *et al*., 2015). Knockout or strong knockdown of single TPC subunits results in pollen and seedling lethality (Gadeyne *et al*., 2014; Van Damme *et al*, 2006; Wang *et al*, 2019). Mild knockdown of TPC subunits or destabilization of TPC by mutating the evolutionary most conserved domain (the WDX domain) in the TPLATE subunit, however, results in viable plants, allowing to address possible developmental functions for this complex (Bashline *et al*, 2015; Van Damme *et al*., 2006; Wang *et al*, 2021).

In this study, we took advantage of WDX domain-dependent TPC destabilization to explore how reduced TPC-dependent endocytic capacity affects receptor-mediated signaling in plants. We compared the response of control plants (*tplate(-/-)* complemented with TPLATE-GFP) with that of plants expressing the partially functional allele (*tplate(-/-)* complemented with WDXM2-GFP) upon exposure to different types of exogenous peptides.

## Results and discussion

### Reduced TPC-dependent endocytosis confers hypersensitivity to a subset of CLE peptides

*In vitro* bioassays comparing root growth in the presence or absence of exogenous peptide ligands provide an easy readout and are widely employed to evaluate how plants respond to peptide-dependent signaling (Anne *et al*, 2018; Blumke *et al*, 2021; Breda *et al*, 2019; Graeff *et al*, 2020; Hazak *et al*, 2017; Hu *et al*, 2018; Poncini *et al*, 2017). To correlate peptide-dependent receptor signalling with CME capacity, we selected several classes of peptide ligands. CME has been shown to internalize the pattern recognition receptors PEP RECEPTOR1 (PEPR1) and FLAGELLIN SENSING 2 (FLS2), which are the respective receptors of the *Arabidopsis thaliana* endogenous elicitor peptides (AtPEPs) and the bacterial peptide FLAGELLIN 22 (FLG22) (Mbengue M, 2016; Ortiz-Morea *et al*., 2016). We also included the C-TERMINALLY ENCODED PEPTIDE 5 (CEP5), which impacts on primary root length and lateral root initiation via its proposed receptor XYLEM INTERMIXED WITH PHLOEM 1 (XIP1)/CEP RECEPTOR 1 (CEPR1) (Roberts *et al*, 2016). Finally, we included fourteen CLV3/EMBRYO SURROUNDING REGION (CLE) peptides, which are essential for shoot and root meristem maintenance by activating various plasma membrane-bound receptors (Yamaguchi *et al*., 2016).

TPLATE and WDXM2 complemented seedlings were grown in the presence of different CLE peptides. The majority of the tested CLE peptides, which were applied at nanomolar concentrations, elicited a similar response in WDXM2 and TPLATE seedlings (Fig EV1A-B). However, we observed a strong hypersensitivity of WDXM2 seedlings to CLV3, CLE10 and CLE40 (Fig EV1A-B). CLE40 is the closest homolog of CLV3 in *Arabidopsis*, and both peptides are crucial for root and shoot meristem maintenance (Brand *et al*, 2000; Clark *et al*, 1995; Fletcher *et al*, 1999; Hobe *et al*, 2003; Ito *et al*, 2006; Schlegel *et al*, 2021; Stahl *et al*., 2013; Stahl *et al*, 2009; Yamaguchi *et al*., 2016). The specific hypersensitivity of the WDXM2 expressing seedlings to these two closely related peptides hinted towards a connection between TPC-dependent endocytosis and CLV1-type receptor signalling.

We subsequently treated TPLATE and WDXM2 complemented plants with CLV3, CLE40 at a concentration of 10 nM as well as with different doses of FLG22, AtPEP1 and CEP5 peptides, previously shown to affect root growth (Poncini *et al*., 2017), and we compared the effect between our two backgrounds that differ in their endocytic capacity (Wang *et al*., 2021). After a 5-day exposure, both WDXM2 and TPLATE seedlings, grown in the presence of the peptides, showed reduced root growth compared to the control situation, indicating that they responded to the treatments. In contrast to the clearly differential effect observed for CLV3 and CLE40 (Fig 1A-B), both backgrounds responded similarly to FLG22 treatment and only a slight but statistically significant difference was found in response to the low dose of AtPEP1 but not to the higher dose (Fig 1C-D). We also did not observe any differential response between TPLATE and WDXM2 complemented plants to both low and high doses of CEP5, although the latter severely reduced root growth (Fig 1C-D). These results indicate that the differential endocytic capacity between both backgrounds elicits hypersensitivity to CLE peptides, but that the mild endocytic flux difference between both backgrounds is insufficient to generate a differential developmental effect due to FLG22-, AtPEP1- or CEP5-dependent receptor signaling at the concentrations used. We conclude that regulatory mechanisms controlling the activity of those receptors remain sufficiently active in both genetic backgrounds.

**Figure 1.**
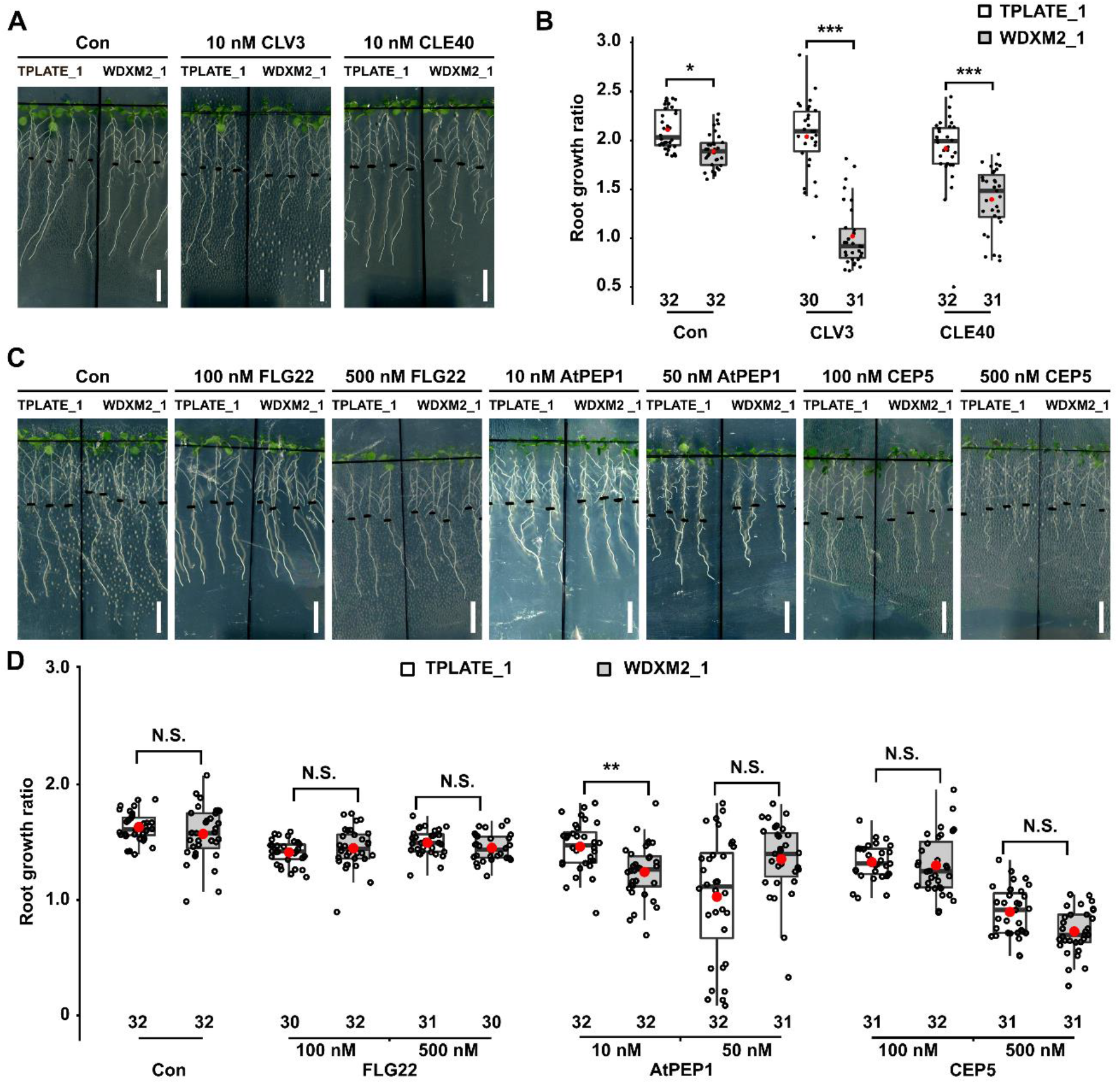
Impaired TPC-dependent endocytic capacity confers hypersensitivity to CLV3 and CLE40 peptides. (A-B) Representative images and quantification of the root growth ratios of TPLATE_1 and WDXM2_1 seedlings (see Table EV1 for the specifications of the lines) treated with or without (Con) low doses of CLV3 or CLE40 peptides. Scale bar = 1cm. (C-D) Representative images and quantification of the root growth ratios of TPLATE_1 and WDXM2_1 seedlings treated with or without (Con) different doses of FLG22, CEP5 and AtPEP1 peptides. 5-day-old seedlings grown vertically on ½ MS medium plate were transferred to freshly prepared ½ MS medium plates supplemented with or without low doses of peptides and grown vertically for an extra 5 days. For each individual root, the primary root length after the transfer was divided by the root length of the seedling before the transfer. The numbers at the bottom of the box plot and jitter box graphs represent the number of individual roots measured. The box plot extends from the 25th to 75th percentiles. The line inside the box marks the median. The whiskers go down and up to the 95% percentile. Data information in panel (B) and (D): Differences as compared to TPLATE complemented lines are indicated (selected pairs from Welch’s ANOVA post hoc pairwise comparison with the Tukey contrasts); N.S., no significant difference; *P < 0.05; **P < 0.01; ***P < 0.001. The data represented results from at least 4 sets of seedlings grown on separate plates.

To independently confirm the observed hypersensitivity to CLV3 and CLE40, we tested the *twd40-2-3* mutant. This is a mild knockdown allele of the TPC subunit TWD40-2 (Bashline *et al*., 2015). Similar to our partially functional WDXM2 allele, *twd40-2-3* mutant plants also exhibited a hypersensitive response to low doses of CLV3 and CLE40 treatment (Fig EV2). Altogether, these results revealed that reduced TPC-dependent endocytosis enhances CLV3 and CLE40 signalling in Arabidopsis roots.

### TPC-dependent endocytosis contributes to SAM maintenance through the WUSCHEL signalling pathway

Next to root meristem maintenance, CLV3-dependent signaling is also essential to maintain SAM homeostasis. Long-term synthetic CLV3 peptide treatment dampens cell proliferation and thus consumes SAM (Hu *et al*., 2018; Ishida *et al*, 2014). To investigate the importance of TPC-dependent endocytosis for SAM maintenance, we compared the sensitivity of TPLATE and WDXM2 complemented plants to long-term CLV3 peptide treatment. Seedling morphologies indicated that TPLATE and WDXM2 seedlings were equally capable of maintaining their SAM in the presence of very low doses of exogenous CLV3 peptides (10 nM), even during long-term treatment (Fig 2A-B). However, higher concentrations (100 nM and 1 μM) of CLV3 revealed hypersensitivity of WDXM2 seedlings and increasingly caused SAM termination in independent mutant WDXM2 lines (Fig 2A-B and Fig EV3A-B). The hypersensitivity of WDXM2 plants to CLV3 further correlated with the protein levels of the complementation constructs in the complemented *tplate* mutant lines (Fig EV3C). These results suggest that TPC-dependent endocytic deficiency causes a dose-dependent hypersensitivity to CLV3-dependent receptor signalling.

**Figure 2.**
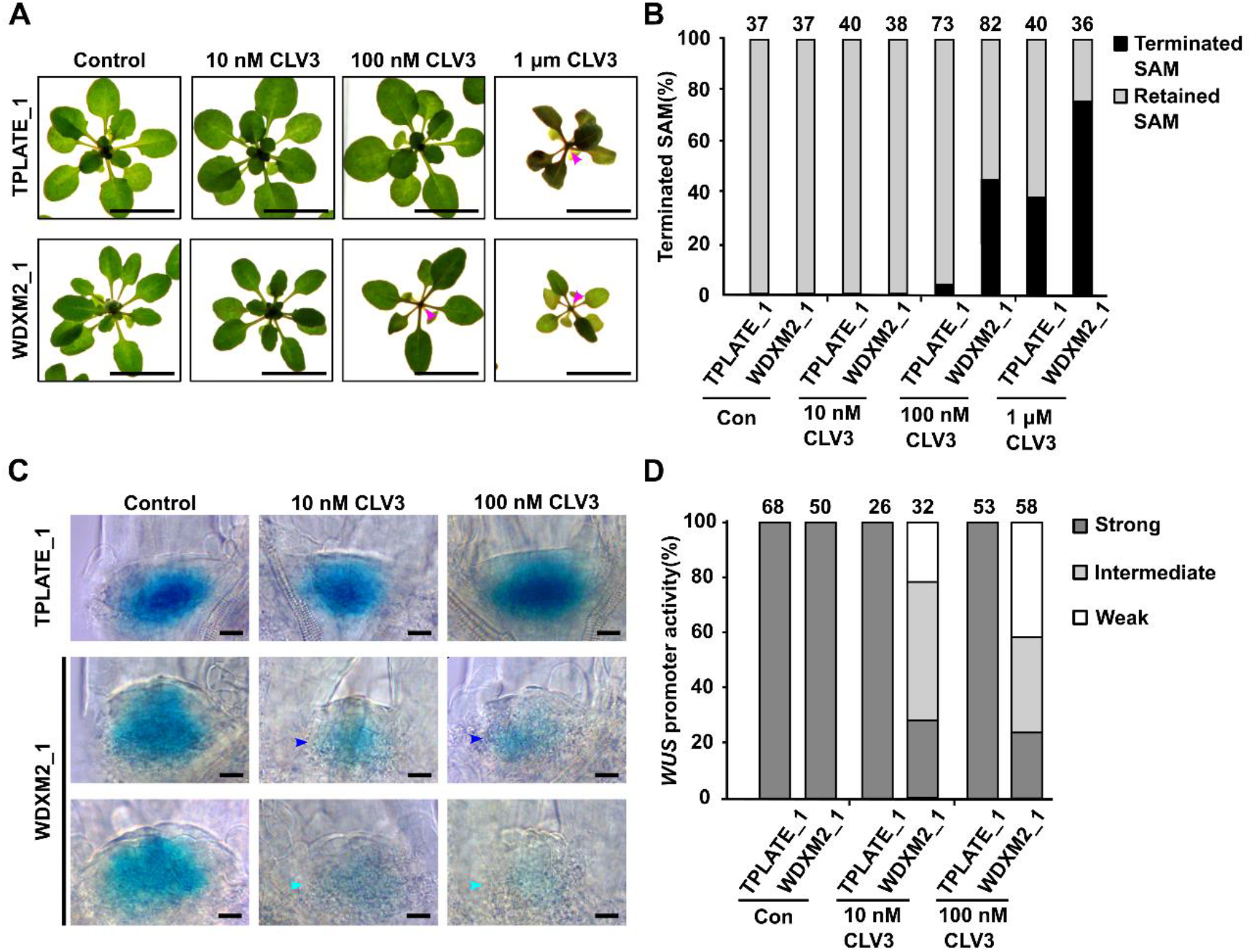
Impaired TPC-dependent endocytic capacity confers hypersensitivity to CLV3 in SAM. (A) Phenotypic comparison of 3- to 4-week-old TPLATE_1 and WDXM2_1 rosette stage plants grown on ½ MS with or without different doses of CLV3 peptide. Magenta arrows indicate terminated SAMs. Scale bar = 1cm. (B) Quantification of the amount of terminated shoot apical meristems in relation to the dose of CLV3 applied. The number of plants used for the quantification is indicated at the top of the bar chart. (C-D) Representative images (C) and quantification (D) of *WUS::GUS* expression in the vegetative SAMs of 3-day-old TPLATE_1 and WDXM2_1 seedlings treated with or without different doses of CLV3 peptide. Intermediate (blue arrowhead) and weak (cyan arrowhead) *WUS* expression is indicated in the SAMs of WDXM2_1 seedlings after CLV3 treatment. Scale bar = 50 µm. *WUS* expression after CLV3 treatment was visually scored and quantified. The numbers of seedlings analyzed is indicated at the top of the bar chart. The data represented in panel A results from at least 5 sets of seedlings grown on separate plates. The data represented in panel D is the combination of two independent repetitions.

In the SAM, CLV3 signalling functions in a negative feedback circuit to dampen stem cell proliferation by regulating the expression of the homeodomain transcription factor WUSCHEL (WUS) (Hazak & Hardtke, 2016; Kitagawa & Jackson, 2019; Yamaguchi *et al*., 2016). To further examine whether TPC-dependent endocytosis is involved in the CLV–WUS feedback loop to regulate SAM homeostasis, we analyzed the expression patterns of *WUS* in TPLATE and WDXM2 complemented plants following a three-day CLV3 peptide treatment. Both 10 nM and 100 nM CLV3 peptide treatment did not visibly impair *WUS* promoter activity in TPLATE vegetative SAMs at the seedling level compared to control conditions as visualized by GUS staining (Fig 2C-D). In WDXM2 vegetative SAMs, however, CLV3 application dampened *WUS* expression in a dose-dependent manner (Fig 2C-D), which is coherent with the terminated SAM phenotype observed at the rosette stage level upon prolonged treatment (Fig 2A-B). These findings reveal that TPC-dependent endocytosis is involved in the regulation of CLV3-WUS signaling in the SAM.

### TPC-dependent endocytosis internalizes CLV1 to dampen CLV3-dependent signalling

The receptor kinase CLV1 signals in response to CLV3 and plays a central role in shoot meristem maintenance (Brand *et al*., 2000; Clark *et al*., 1997; Fletcher *et al*., 1999; Ogawa *et al*, 2008; Shinohara & Matsubayashi, 2015; Somssich *et al*, 2015). CLV1 levels increase at PM in the absence of CLV3 and accumulate in the vacuole in the presence of CLV3 (Nimchuk *et al*., 2011). CLV3-induced vacuolar accumulation of CLV1 suggests a negative regulation of CLV3/CLV1 signaling by internalization, yet this hypothesis remains to be experimentally tested (Yamaguchi *et al*., 2016).

To characterize whether TPC-dependent endocytosis functions in CLV1 internalization, we evaluated whether the response of TPLATE and WDXM2 complemented plants to CLV3 treatment depended on the presence of CLV1. Combining the *clv1-101* null allele (Atsuko Kinoshita, 2010) with our TPLATE and WDXM2 complemented plants largely suppressed the hypersensitivity to exogenous CLV3 leading to SAM termination in WDXM2, although not completely (Fig 3A-B). Combining the strong and dominant-negative *clv1* mutant allele *clv1-8* (Clark *et al*., 1997; Dievart *et al*., 2003) restored SAM maintenance in WDXM2 in the presence of 100nM CLV3 (Fig 3A-B). The differential effect of exogenous CLV3 on SAM activity between WDXM2, WDXM2/*clv1-101* and WDXM2/*clv1-8* was also apparent in the number of leaves that the plants produced (Fig 3C).

**Figure 3.**
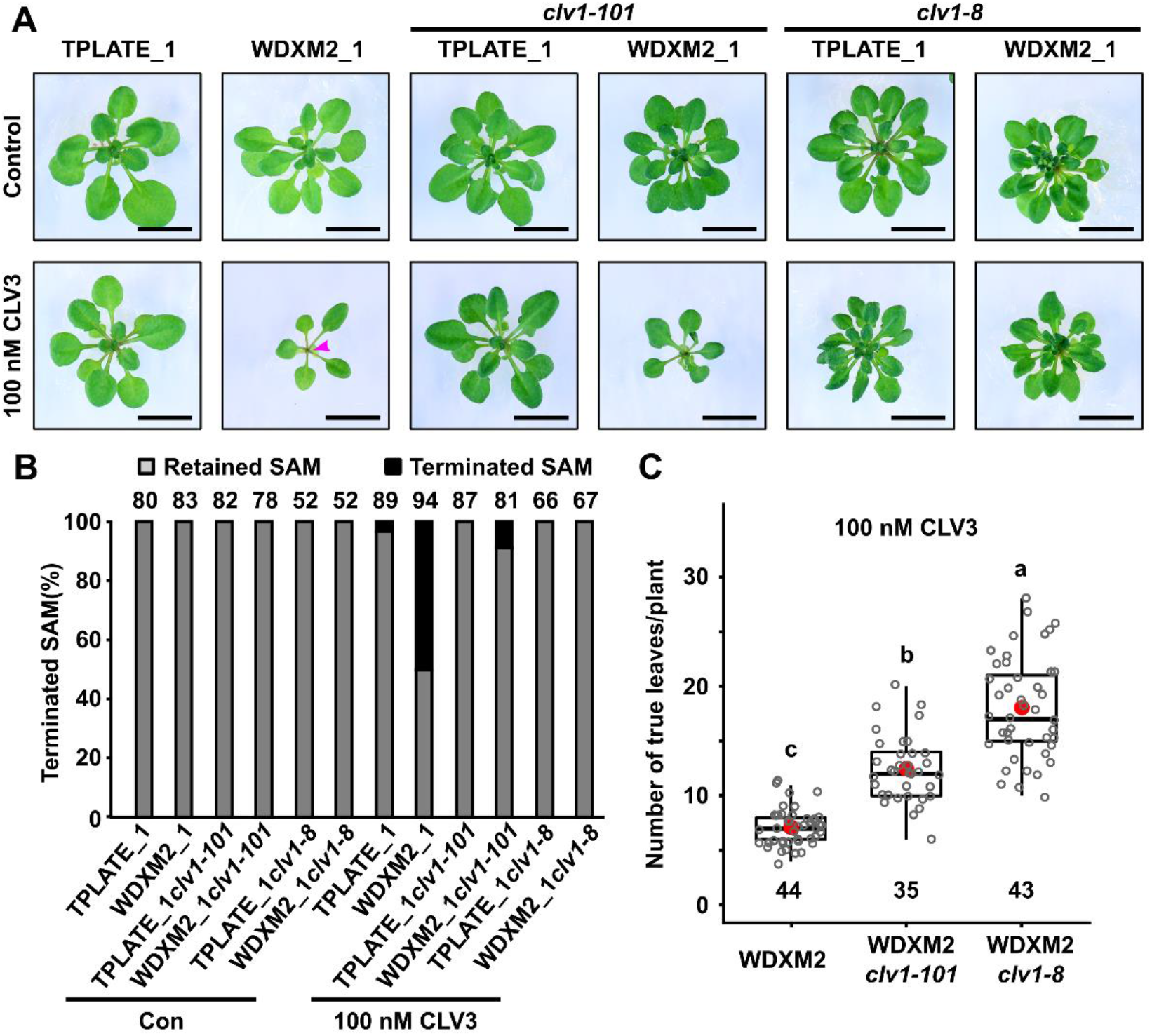
CLV1 loss-of-function dampens CLV3 hypersensitivity in the SAMs of WDXM2 complemented plants. (A) Phenotypic comparisons of 3 to 4 -week-old TPLATE_1 and WDXM2_1 plants as well as combinations of these with the *clv1* null (*clv1-101*) or dominant negative (*clv1-8*) mutant backgrounds under control conditions or in the presence of 100 nM exogenous CLV3 peptide. The magenta arrowhead indicates a terminated SAM. Scale bar = 1cm. (B) Quantification of the amount of terminated meristems in relation to the dose of CLV3 peptide applied. Numbers of plants used for quantification are indicated at the top of the bar chart. (C) Box plot and jitter box representation of the quantification of the number of leaves produced by WDXM2, WDXM2/*clv1-101* and WDXM2/*clv1-8* plants grown *in vitro* on medium supplemented with 100nM CLV3. Numbers of biological samples are indicated at the bottom of the box plot and jitter box graphs. The box plot extends from the 25th to 75th percentiles. The line inside the box marks the median. The whiskers go down and up to the 95% percentile. Letters (a, b and c) represent significantly different groups (P < 0.001) evaluated by Welch’s ANOVA post hoc pairwise comparison with the Tukey contrasts. The data represented in panel B results from at least 6 sets of seedlings grown on separate plates. The data in panel C is based on a random selection of 3 to 4 plates from panel B.

These results reveal that CLV1 predominantly contributes to the hypersensitivity of CLV3-dependent signalling in WDXM2 mutant plants. The different capacity of the *clv1-101* null allele and the *clv1-8* dominant negative allele to reduce the sensitivity of WDXM2 to CLV3 is likely attributed to genetic redundancy within the CLV1 receptor family (DeYoung *et al*., 2006; Deyoung & Clark, 2008; Nimchuk, 2017; Nimchuk *et al*, 2015; Shinohara & Matsubayashi, 2015).

The WDXM mutation destabilizes TPC and thereby negatively affects endocytic capacity (Wang et al., 2021). The entire TPC complex is required to execute CME at PM (Gadeyne *et al*., 2014; Johnson *et al*, 2021; Wang *et al*, 2020; Wang *et al*., 2021; Yperman *et al*, 2021b) and destabilizing TPC in WDXM2 complemented plants impairs endocytic capacity while it does not affect recruitment of the two AtEH/Pan1 subunits at PM, which are involved in promoting autophagy (Wang *et al*., 2021; Wang *et al*., 2019). It is therefore likely that the CLV1-dependent hypersensitivity to CLV3 is linked to altered endocytosis of CLV1 in WDXM2. CLV1 is a master regulator of flower development (Clark *et al*., 1997; Schoof *et al*, 2000). Both TPLATE and WDXM2 are expressed in the inflorescence meristem at roughly similar levels although in these tissues, WDXM2 appears to be slightly less PM-associated compared to TPLATE (Fig EV4A). Next, we monitored the PM localization of functional CLV1-GFP in the inflorescence meristems of TPLATE and WDXM2 plants. We observed similar levels of CLV1-GFP on the PM in WDXM2 inflorescence meristems compared with TPLATE plants (Fig EV4). In vegetative meristems, however, our live imaging analysis clearly showed increased levels of CLV1-GFP in the WDXM2 background (Fig 4A).

**Figure 4.**
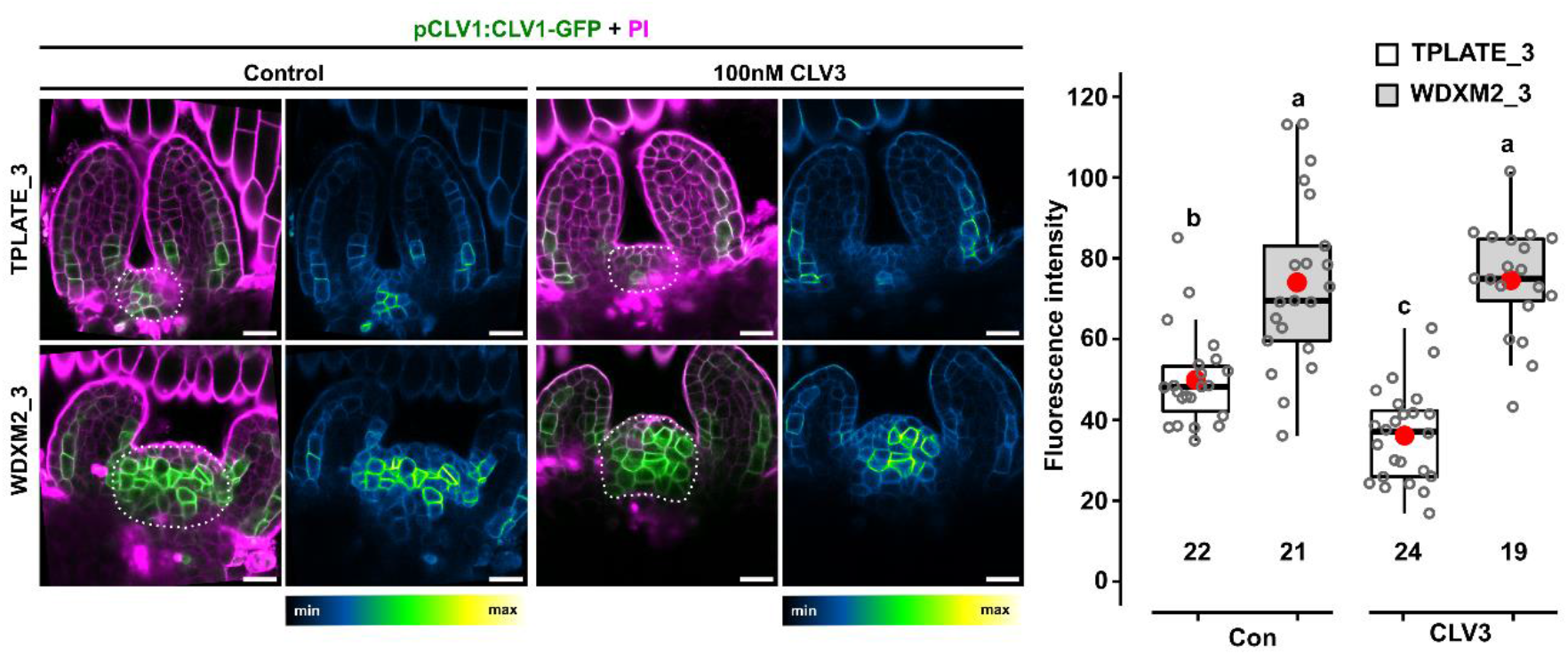
Reduced TPC-dependent endocytic capacity impairs internalization of CLV1 from the PM in SAM cells. Confocal images and quantification of Arabidopsis seedlings showing enhanced PM localization of CLV1-GFP in WDXM2_3 vegetative meristems compared to vegetative meristems of TPLATE_3 lines with or without exogenous CLV3 (100 nM) in the growing medium from germination onward. Left panels are merged channels (GFP and PI), right panels are GFP-only channels represented via an intensity scale. Scale bar = 20 µm. The Box plot and jitter box representation graph shows the average fluorescence intensity (8-bit gray values) of CLV1 over the entire SAM (indicated by a dotted line). The box plot extends from the 25th to 75th percentiles. The line inside the box marks the median. The whiskers go down and up to the 95% percentile. Numbers of biological samples from two repeats are indicated at the bottom of the box plot and jitter box graphs. Differences of CLV1-GFP intensity between WDXM2_3 and TPLATE_3 lines under both conditions were evaluated by Welch’s ANOVA post hoc pairwise comparison with the Tukey contrasts. Letters (a, b and c) represent significant difference between groups (a,b,c; P < 0.001). The quantification is a combination of two independent experiments for each genotype and treatment.

CLV1 undergoes CLV3-mediated degradation in inflorescence meristems upon induction of *CLV3* expression in the *clv3-2* mutant background (Nimchuk et al., 2011). In vegetative meristems and in the presence of endogenous levels of CLV3, signal intensities of CLV1 showed variation before and after exogenous CLV3 application. Live cell imaging of the same vegetative meristem before and after CLV3 addition (Fig EV5A) as well as quantification of treated and untreated meristems however revealed that CLV1 levels significantly reduced upon long-term (present in the medium from germination onward; Fig 4) or short-term (10 and 30min; Fig EV5B-E) exogenous CLV3 application in TPLATE seedlings, while this was not the case in WDXM2 seedlings (Fig 4; Fig EV5B-E).

These results strongly correlate the endocytosis deficiency in WDXM2 with impaired internalization of CLV1 in vegetative meristems. Increased CLV1 levels at PM are also in accordance with the fact that WDXM2 complemented plants are hypersensitive to CLV3 peptide treatment, which correlates with strongly reduced *WUS* levels and therefore likely increased CLV1-mediated transcriptional repression (Fig 2 and Fig 3). Despite this hypersensitivity, vegetative SAMs in WDXM2 appear enlarged compared to those in TPLATE control seedlings (Fig 4 and Fig EV5). How this relates to the abundance of CLV1 at PM and to altered *WUS* levels remains to be determined.

To establish a direct link between CLV1 and TPC, we examined the interaction between TPC and CLV1. TPC, visualized using an antibody against TPLATE, specifically co-purified with CLV1 in Arabidopsis seedlings when CLV1-2xGFP was used as bait (Fig 5A). Next, we aimed to confirm this interaction and to determine which adaptor complex subunits were involved. Tyrosine motif-based cargo recognition involves the medium subunit of the Adaptor protein 2 complex, AP-2M (Arora & Damme, 2021), whose counterpart in TPC is the TML subunit. Furthermore, TPLATE co-purified with CLV1 (Fig 5A) and AtEH1/Pan1 was shown to interact with cargo (Yperman *et al*, 2021a). We therefore selected these proteins for ratiometric bimolecular fluorescence complementation (rBiFC) in *N. benthamiana*. Similar to previous experiments, the shaggy-like kinase BIN2 served as negative control (Arora *et al*, 2020). We could not visualize interaction between CLV1 and TPLATE, TML or AP-2M in this system (Fig 5B-C). Our confocal analysis, however, clearly linked CLV1 to the plant-specific TPC subunit AtEH1/Pan1 (Gadeyne *et al*., 2014; Hirst *et al*, 2014) in the presence and absence of exogenous CLV3 peptide (Fig 5B-C). The interaction between CLV1 and AtEH1/Pan1 was further assessed via yeast-two-hybrid (Y2H) using the cytoplasmic part of CLV1 and the N-terminal part of AtEH1/Pan1 ending just after the second EH domain (Yperman *et al*., 2021b). In total, 24 independent double transformations, combining CLV1 with AtEH1/Pan1, CLV1 with empty vector control or AtEH1/Pan1 with empty vector control were compared, alongside 8 double transformations of the empty vector control and the p53-SV40 positive control (Fig 5D). The results clearly show a specific interaction between CLV1 and AtEH1/Pan1 (Fig 5D). Both rBiFC and Y2H therefore clearly link the cytoplasmic part of CLV1 to the N-terminal part of AtEH1/Pan1. The N-terminal located EH domains of AtEH1/Pan1 were previously also shown to be involved in membrane recruitment of TPC as well as in the internalization of the Secretory Carrier Membrane Protein 5 (SCAMP5) via its double NPF motif (Johnson *et al*., 2021; Yperman *et al*., 2021a). CLV1, in contrast to SCAMP5, does however not contain obvious NPF motifs. How CLV1 is recognized by AtEH1/Pan1 therefore remains to be determined.

**Figure 5.**
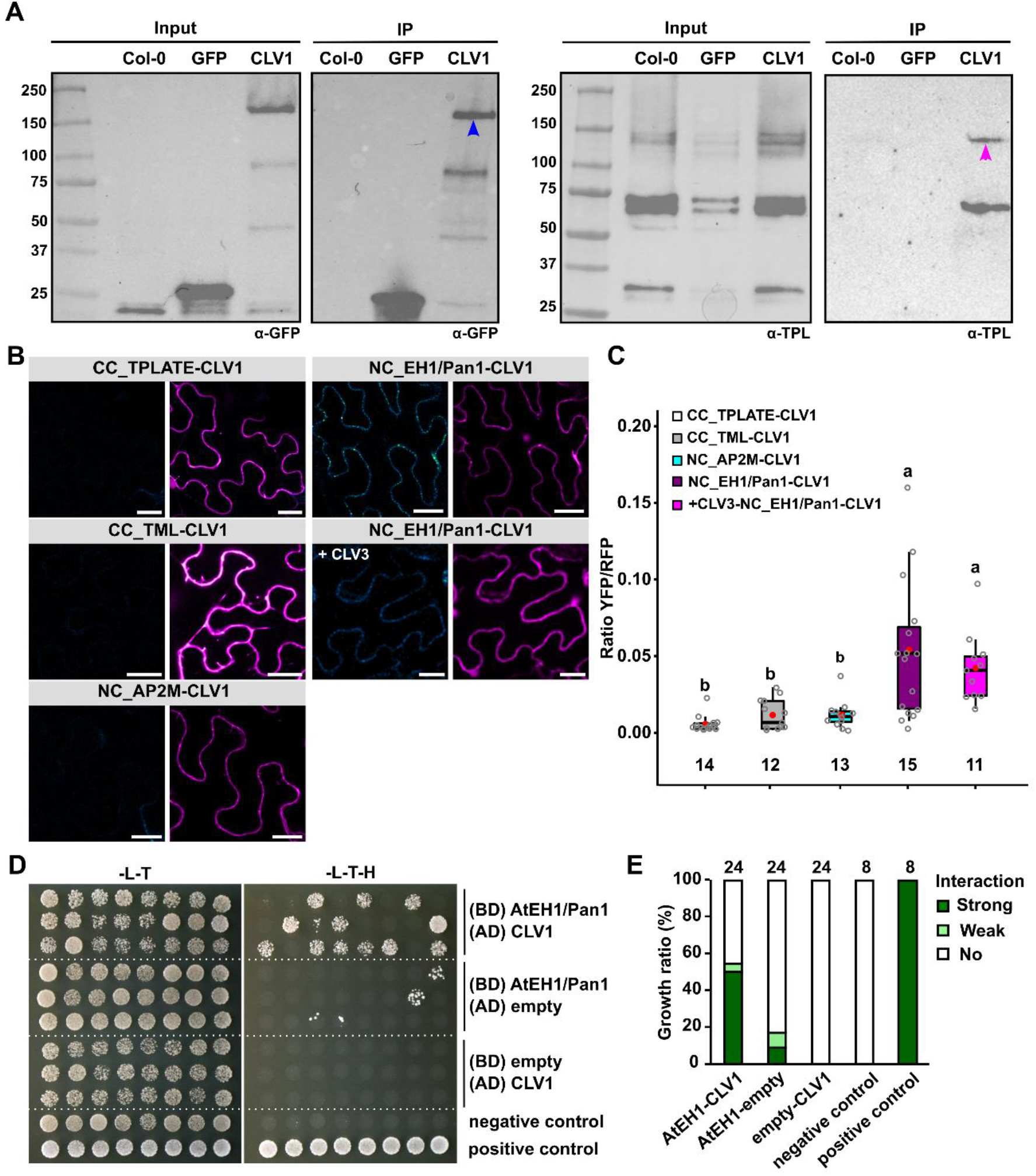
TPC interacts with CLV1 through its AtEH1/Pan1 subunit. (A) Co-immunoprecipitation experiment comparing WT (Col-0) Arabidopsis seedlings and seedlings expressing pCLV1::CLV1-2xGFP (CLV1) and 35S::eGFP (GFP). CLV1 specifically co-purifies with endogenous TPLATE. The blue arrow marks full length CLV1 and the magenta arrow marks full length TPLATE. Numbers next to the ladder represents the protein molecular weight (kDa). The experiment was independently performed twice with an identical result. (B-C) Representative confocal images and quantification of ratiometric BiFC analyses exploring the interaction between TPC subunits TPLATE, TML and AtEH1/Pan1, the AP-2 complex subunit AP2M, and CLV1. The identified interaction between CLV1 and AtEH1/Pan1 was also performed in the presence of exogenous CLV3 peptide application (1 µM in infiltration buffer). CC and NC refer to the orientation of the nYFP and cYFP halves of YFP fused to both proteins. CLV1 was always tagged C-terminally. Left panels in (C) represent the YFP channel, shown via an intensity scale whereas the right panels represent the RFP control channel (free RFP, magenta) against which the intensity of the YFP BiFC channel was normalized. Scale bars = 25 µm. (C) Box plot and jitter box representation showing the quantification of the YFP/RFP fluorescence ratios from two independent experiments. The box plot extends from the 25th to 75th percentiles. The line inside the box marks the median. The whiskers go down and up to the 95% percentile. Numbers of biological samples from at least two independent transformations are indicated at the bottom of the graph. Letters (a, b and c) represent significantly different groups (P < 0.001) evaluated by Welch’s ANOVA post hoc pairwise comparison with the Tukey contrasts. (D-E) Yeast two hybrid analysis (D) and respective quantification (E) between the cytoplasmic part of CLV1 (AA 671-980) and the N-terminal part of AtEH1/Pan1, which ends after the second EH domain (AA 1-527). Combining CLV1 in pGADT7 (AD) with AtEH1/Pan1 in pGBKT7 (BD) allowed growth on selective medium (-L-T-H; strong in 12/24 and weak in 1/24 independent double transformations) whereas only 2/24 transformations showed strong and 2/24 showed weak growth on selective medium in the controls, likely caused by some level of auto-activation of AtEH1/Pan1. The negative control consisted of both empty pGBKT7 and pGADT7 vectors (8 independent double transformations) and the positive control (eight independent double transformations) combined pGADT7-SV40 T-Ag with pGBKT7-p53. No: no growth observed on -L-T-H. The data shown represents individual double transformants and the assay was technically repeated twice.

Taken together, our findings reveal that the hypersensitivity of WDXM2 complemented plants to CLV3 is most likely a consequence of sustained signalling from the PM in vegetative meristems, which is caused by impaired internalization of CLV1 due to reduced TPC-dependent endocytosis. TPC-dependent endocytosis, therefore, serves to internalize CLV1 to attenuate CLV3 signalling to prevent meristem termination. Our work thus identifies CME as a mechanism to control the availability of CLV1 at the PM and to tune the activity of the shoot stem cell niche during plant development.

## Materials and Methods

### Molecular cloning

mSCARLET (Bindels *et al*, 2017) was amplified with a stop codon from plasmid pEB2-mSCARLET (Addgene,104006), introduced into pDONRP2R-P3 via a Gateway BP reaction (Invitrogen) and confirmed by sequencing. To generate mSCARLET-fused expression constructs of TPLATE and WDXM2, The pDONR221-TPLATE and pDONR221-WDXM2 motif substituted entry clones (Wang *et al*., 2021) were combined with pHm34GW (Karimi *et al*, 2007), pDONRP4-P1r-Lat52 (Van Damme *et al*., 2006), and pDONRP2R-P3-mSCARLET in triple gateway LR reactions (Invitrogen).

The pBiFCt-2in1 BiFC vectors, which allow quantification of the observed Bimolecular YPF fluorescence complementation by measuring the ratio between the intensity of the YPF signal for a specific pair of interacting proteins and the intensity of the constitutively expressed RFP which is present on the backbone of the vector, were used to generate CLV1 related rBiFC constructs (Grefen & Blatt, 2012). The CLV1 entry clone for rBiFC reactions was amplified from a published plasmid (Schlegel *et al*., 2021), while TPLATE, TML, AtEH1/Pan1 and AP2M were obtained from previously reported rBiFC experiments (Arora *et al*., 2020; Liu *et al*, 2020; Yperman *et al*., 2021a). Entry clones were assembled in an empty rBiFC destination vector (pBiFCt-2in1-CC, Addgene 105114 or pBiFCt-2in1-NC, Addgene 105112) with a Gateway LR recombination reaction and selected using LB containing spectinomycin and XgalI. The final rBIFC vectors were checked by restriction digestion and sequencing of the recombination borders. For Y2H, the N-terminal domain of AtEH1/Pan1 (AA 1-527) was amplified using following primer pairs (AtEH1_1-527_GBD_F GCCATGGAGGCCGAATTCCCAATGGCGGGTCAGAATCCTAACATGG and AtEH1_1-527_GBD_R CTGCAGGTCGACGGATCCCCTTATGCAGAATATCCATTACCTAGGTGATTAGC) and cloned into the pGBKT7 vector (Clontech). The cytoplasmic part of CLV1 (AA 671 to 980, corresponding to the end of the transmembrane helix, from amino acids LAWKL to the end of the uniprot sequence Q9SYQ8) was amplified using following primer pairs (CLV1_671-980_GAD_F GAGGCCAGTGAATTCCACCCACTCGCCTGGAAACTAACCGCCTTC and CLV1_671-980_GAD_R TCCCGTATCGATGCCCACCCTTAGAACGCGATCAAGTTCGCCACGG) and cloned into pGADT7 (Clontech). Both vectors were generated via Gibson assembly following SmaI-dependent linearization of the vectors. Plasmids were verified using sequencing.

### Arabidopsis transgenic lines and growth conditions

All plant materials used in this research are in the Columbia-0 (Col-0) ecotype background. Information on plant materials is listed in Table EV1. To generate the mSCARLET fusions of transgenic lines, *tplate* heterozygous mutant plants were identified by genotyping PCR and were transformed with expression constructs of TPLATE and WDXM2 fused to mSCARLET under the control of LAT52 promoter as described before (Van Damme *et al*., 2006; Wang *et al*., 2021; Yperman *et al*., 2021b). Primary transformants were selected with Hygromycin, and those carrying the *tplate* T-DNA insertion were identified via genotyping PCR. The complemented lines in the T2 generation were further genotyped to identify homozygous *tplate* mutants (Wang *et al*., 2021).

For all the crosses, the same reporter line or mutant plant was used as male to cross with TPLATE and WDXM2 complemented lines respectively. The *pWUS::GUS* (Su *et al*, 2009) reporter line was crossed into TPLATE_1 and WDXM2_1 complemented mutant backgrounds. In the progeny, F2 plants were genotyped to obtain homozygous *tplate* mutant backgrounds. The F3 or F4 generation plants were screened to identify homozygous plants for *pWUS::GUS* expression by GUS staining. The *clv1* null mutant *clv1-101* (Atsuko Kinoshita, 2010) and the dominant-negative *clv1-8* mutant (Dievart *et al*., 2003) were crossed into the TPLATE_1 and WDXM2_1 complemented lines. The F2 or F3 generation plants were genotyped or sequenced to identify the *tplate/clv1-101 or tplate/clv1-8* double mutant backgrounds. To introduce the CLV1 marker line into TPLATE and WDXM2 complemented lines, a wild type Col-0 plant expressing the functional pCLV1::CLV1-GFP (Schlegel *et al*., 2021) was backcrossed to Col-0 and a single locus expression F2 line was identified by segregation using Basta (20 mg/L) selection. Then the F2 Basta resistant CLV1-GFP expressing plant was used to cross with the TPLATE_3 and WDXM2_3 complemented plants. In the progeny, plants homozygous for the *tplate* mutant background were identified by genotyping PCR while homozygous expression of CLV1-GFP was selected by segregation on BASTA. For co-IP experiments, the pCLV1::CLV1-2xGFP line was used (Nimchuk *et al*., 2011).

Seeds were sterilized by chlorine gas sterilization and sown on ½ MS medium plates without sugar following a 3-day vernalization period at 4°C. Seedlings were grown in a growth chamber under continuous light conditions at 21°C.

### Phenotypic analysis

Sequences of CLE peptides described before (Yamaguchi *et al*., 2016) were ordered from GeneScript. Information on the peptides is listed in Table EV2. For shoot treatments, seedlings were grown horizontally on ½ MS medium supplemented with or without the indicated concentration of CLV3 peptide for 3 weeks. Plants with terminated shoots were counted manually. For root growth assays, seedlings were initially grown on ½ MS medium supplemented with or without CLE peptides for a certain duration (data depicted in Figure EV1). For the FLG22, AtPEP, CEB5, CLV3 and CLE40 peptides depicted in Figure 1 and in Figure 2, seedlings were grown on ½ MS plates and then transferred to plates with and without the indicated amount of peptides. Plates with seedlings were scanned and root lengths were measured with the Fiji software package (https://imagej.net/software/fiji/) equipped with the NeuronJ plugin (Meijering *et al*, 2004). Quantification of the number of leaves in Figure 3C was done manually using the cell counter plugin in Fiji.

### GUS staining

GUS staining was performed as described before (Lammens *et al*, 2008). Seedlings (3-days after putting the plates in continuous light, i.e. roughly one day after germination) expressing *pWUS::GUS* grown on ½ MS with or without CLV3 peptide were harvested and incubated with 80% cold acetone for 30 min. After that, seedlings were washed with phosphate buffer (pH = 7.2), incubated in GUS staining solution (1 mg/ml of 5-bromo-4-chromo-3-indolyl β-D-glucuronide, 2 mM ferricyanide, and 0.5 mM ferrocyanide in 100 mM phosphate buffer pH 7.2) and kept at 37 °C in the dark for 3 hours. After GUS staining, seedlings were cleared with lactic acid and visualized between slide and coverslip on a BX51 light microscope (Olympus) using a 10x or 20x magnification.

### Nicotiana benthamiana infiltration

Three- to four-week-old *Nicotiana benthamiana* plants grown in greenhouse under long-day conditions (06-22 h light, 100 PAR, 21°C) were used for infiltration as described before (Arora *et al*., 2020). 3 days after infiltration, *N. benthamiana* leaves were imaged with an SP8X confocal microscope. CLV3 peptide (1 µM) in infiltration buffer (10 mM MgCl_2_ and 10 mM MES, pH 5.6) was applied via leaf infiltration. After 5 min incubation, the injected samples were imaged within 30 min.

### Live-Cell Imaging and Analysis

A Leica SP8X confocal microscope equipped with a white laser was used for all confocal imaging via a 40x (HC PL APO CS2, NA=1.10) water-immersion corrected objective except the flower meristem imaging.

rBiFC images were acquired with Hybrid detectors (HyDTM) using a time-gated window between 0.3 ns-6.0 ns and in line sequential mode. YFP signals were acquired using WLL 514 nm excitation and an emission window of 520-550 nm, and RFP signals were detected using WLL 561 nm excitation and an emission window of 580-650 nm. All images were taken using the same settings for YFP and RFP detection and saturation was avoided in order not to interfere with the ratiometric quantification.

For CLV1-GFP imaging in vegetative SAMs in Figure 4 and Figure EV5B-E, seeds expressing CLV1-GFP in TPLATE and WDXM2 complemented mutant backgrounds were germinated on ½ MS plates supplemented with or without 100 nM of CLV3 peptide. Seedlings were imaged following 3-days after putting the plates in continuous light, which roughly equals 1 day after germination. For CLV1-GFP imaging upon short-term CLV3 peptide treatment in Figure EV5, seedlings grown on ½ MS plates (3 days in light) were used. After removal of the cotyledons, seedlings were incubated in ½ MS medium containing 1 µM CLV3 peptide and 0.1% Tween 20 (v/v) for 10 or 30 min, and washed with water shortly 3 times. Prior to imaging, seedlings expressing CLV1-GFP were stained with PI solution (10 μg/mL) for 1 to 2 min. The Hybrid detectors (HyDTM) were employed to image PI (excitation at 561 nm, emission between 600-700 nm) and CLV1-GFP (excitation at 488 nm, emission between 500-540 nm) without (PI) or with (GFP) a time-gated window between 0.3 ns-6.0 ns. To achieve sufficient signal when imaging CLV1-GFP in the vegetative SAMs of TPLATE-3 and WDXM2_3 seedlings, accumulative imaging was used. Images were acquired using 8 times line accumulation and 2 times frame averaging.

For the flower SAM imaging in Figure EV4A, Arabidopsis plants were grown in soil for 4 weeks at 21 °C under long day condition (16 h light : 8 h dark, LED 150 µmol/m^2^/s). Primary inflorescence shoot apical meristems were dissected, mounted in ACM and then stained with 100 µM propidium iodide (PI; Merck) for 5 min prior to imaging (Brunoud *et al*, 2020). Meristems were imaged with a Zeiss LSM 710 spectral microscope using the following settings: GFP (excitation at 488 nm, emission between 510-558 nm) and propidium iodide (excitation 488 nm, emission between 605-650 nm).

For the flower SAM imaging in Figure EV4B, SAMs were imaged using a Zeiss LSM 780 confocal microscope (40× water immersion objective, Zeiss C-PlanApo, NA 1.2). Shoot meristems were manually dissected by cutting of the stem, removing the flowers, and were stained with 1 mg/ml DAPI. GFP was excited with an argon laser at 488 nm and emission was detected between 490-530 nm, and mSCARLET was excited with Diode-pumped solid state (DPSS) lasers at 514 nm and detected between 570-650 nm. DAPI was excited at 405 nm with a laser diode and detected between 410-480 nm.

For CLV1-GFP imaging in vegetative SAMs in Figure EV5A, vegetative shoot apices at 3 DAG were manually dissected under a stereo microscope by removing the leaf primordia. The cell wall was stained with PI. After removal of the leaf primordia, vegetative SAMs were treated with ½ MS medium containing 1 µM CLV3 peptide and 0.1% Tween 20 and imaged at 0 min and 30 min after treatment. Z-stacks of vegetative SAMs were acquired using a Zeiss LSM 780 confocal microscope (40× water immersion objective, Zeiss C-PlanApo, NA 1.2). GFP was excited with an Argon laser at 488 nm and emission was detected using a 490-530 nm window. PI was excited at 561 nm by a DPSS laser and detected using a 590-650 nm window.

The quantification of rBiFC and SAM images was performed using Fiji. For rBiFC, a region of interest (ROI) on PM of the cells was selected and the intensities of YFP and RFP signals were measured. The ratios between YFP and RFP signals per cell were then calculated and plotted. For the quantification of CLV1-GFP in TPLATE and WDXM2 vegetative SAMs, a region of interest (ROI) covering the meristem was defined and the CLV1-GFP signal intensities were measured. Only images with less than 1% saturated pixels were quantified. The histogram function in Fiji was used to generate intensity values (8-bit gray values) for each pixel and the top 10% highest intensity pixels were used to calculate the mean fluorescence intensities using an in-house designed script in Microsoft Excel. Using a selection of the strongest intensity pixels for the calculations omits background noise that otherwise reduces the average fluorescence intensities of the quantifications and follows from the rationale that the fluorescence is linked to the endomembrane system and therefore not continuously present throughout the selected ROI. Similar approaches are also used to calculate ratios of endocytic flux between PM and endosomal compartments (Dejonghe *et al*, 2016; Mishev *et al*, 2018).

### Protein extraction and Western blotting

Arabidopsis seedlings were grown for 5 days on ½ MS medium without sugar under continuous light conditions. Seedlings were harvested, flash-frozen, and grinded in liquid nitrogen. Proteins were extracted in a 1:1 ratio, buffer (ml):seedlings (g), in HB+ buffer, as described before (Van Leene *et al*, 2015). Protein extracts were incubated for 30 min at 4°C on a rotating wheel before spinning down twice at 20,000 g for 20 min. The supernatant concentration was measured using the Bradford Protein Assay (Invitrogen), and equal amounts of proteins were loaded on 4 to 20% gradient gels (Bio-Rad). Gels were transferred to nitrocellulose membranes using the Trans-Blot Turbo system (Bio-Rad). Blots were incubated with α-TPLATE appendage antibodies (rabbit) (Dejonghe *et al*, 2019) and imaged on a ChemiDoc Imaging System (Bio-Rad).

#### Co-immunoprecipitation

Finely ground material was suspended in homogenization extraction buffer [150 mM Tris-HCl, 150 mM NaCl, 0.5 mM EDTA, 10% glycerol, 1 mM sodium molybdate, 1 mM NaF, 10 mM DTT, 1% IGEPAL CA-630 (Sigma-Aldrich,USA) with Complete Ultra EDTA-free Protease Inhibitor Cocktail Tablets (Roche, Switzerland; 1 tablet per 10 mL)]. After 30 min of rotation at 4 °C, cell debris was removed from the samples by centrifugation for 15 min at 2000 g at 4 °C. Supernatant was transferred to a new tube through Miracloth (Millipore Sigma, USA). Then, 50 μL pre-equilibrated GFP-Trap®_MA beads (ChromoTek, Germany) was added into each sample and samples were incubated for 2 h at 4 °C to maximize the protein binding. Afterwards, the beads were washed two times with wash buffer (20 mM Tris-HCl pH 7.5, 150 mM NaCl). Protein was eluted from the beads by adding Laemmli sample buffer (Bio-rad, Laboratories, Inc., USA), Sample Reducing Agent (Invitrogen, USA) and incubating at 70 °C for 10 min.

The proteins were separated on 4–15% SDS-PAGE stain-free protein gel (Bio-Rad Laboratories, Inc., USA), followed by transferring onto a Trans-Blot® Turbo™ Mini PVDF Transfer Packs (Bio-Rad Laboratories, Inc., USA). After blocking with 5% Skim Milk (Difco,USA) for 1 h at room temperature, blots were incubated with α GFP-HRP (ChromoTek, Germany) (1:2000) or α TPLATE2 (rabbit) (Dejonghe et al., 2019) overnight at 4 °C. Imaging was done using Chemiluminescent substrate (Thermo Fisher Scientific, USA) and detected by ChemiDoc™ MP Imaging System (Bio-Rad Laboratories, Inc., USA).

### Yeast two hybrid analysis

The N-terminal part of AtEH1/Pan1 (AA 1-527) up to the coiled coil domain in pGBKT7 and the cytoplasmic part of CLV1 (AA 671-980) in pGADT7 were combined with each other and with empty control plasmids using the Matchmaker™ Gold Yeast Two-Hybrid System (Clontech). The vectors were co-transformed into the Y2Hgold MATa Yeast strain. Auto-activation was tested by co-transforming each vector with the corresponding empty pGADT7 and pGBKT7 vectors. The empty pGADT7 and pGBKT7 were also co-transformed as a negative control and as a positive control, we used the pGADT7-SV40 T-Ag and pGBKT7-p53 supplied with the Matchmaker system (Clontech).

Colonies of double transformed yeasts were first selected on SD -Leu -Trp plates. After 3 days at 30 °C, colonies were picked and grown for 3 days in liquid -Leu –Trp medium at 30 °C 200 rpm. Fully grown cultures were diluted 1/5 in -L-T-H and 10 µl was spotted on SD -Leu -Trp and SD -Leu -Trp -His plates. Pictures were taken after 3 days at 30 °C.

### Statistical analysis

The R package in R studio (www.rstudio.com) was used. Data were tested for normality and heteroscedasticity, after which the multcomp package was used (Herberich *et al*, 2010).

## Expanded View figures

**Figure EV1.**
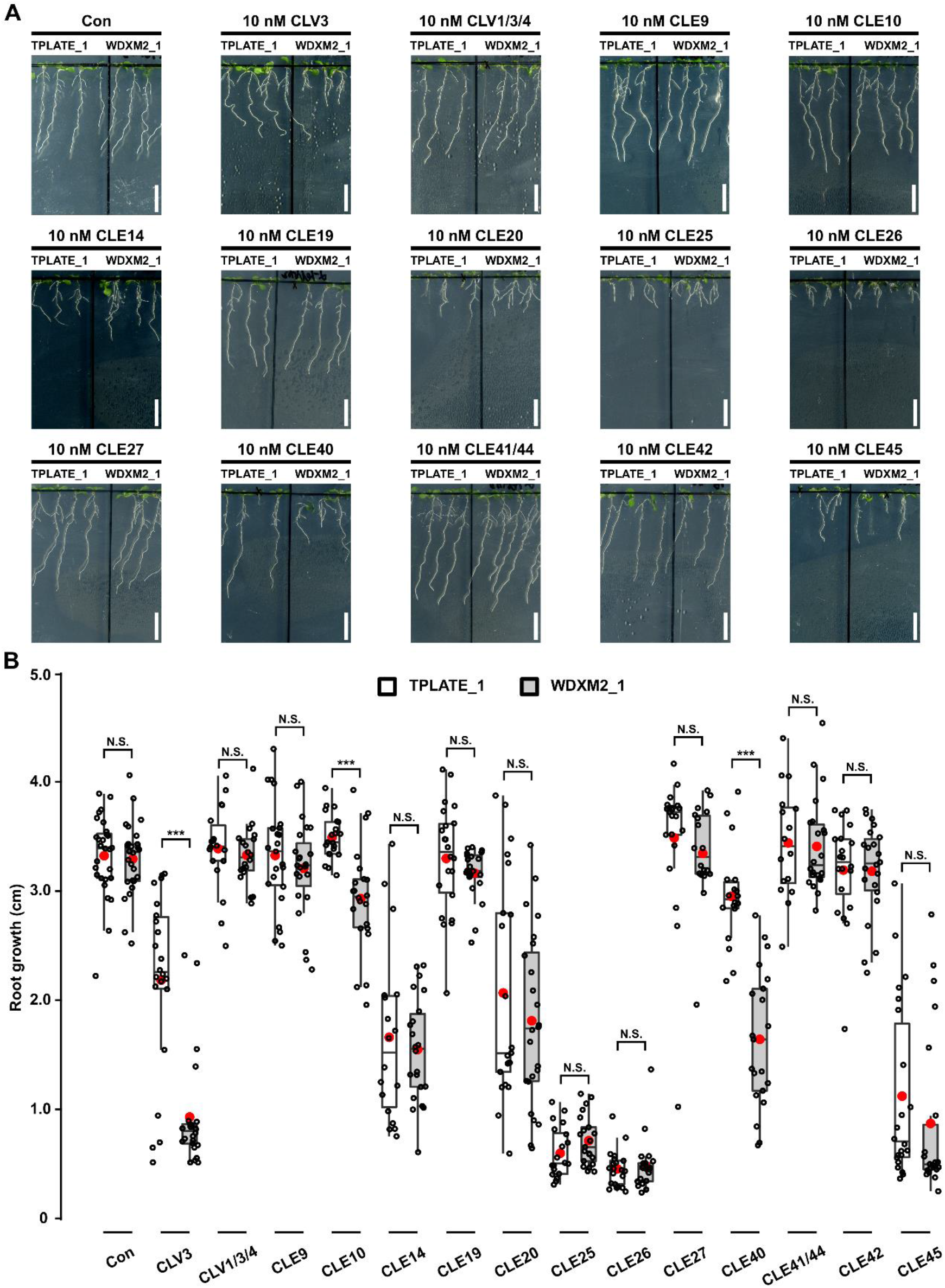
Impaired TPC-dependent endocytic capacity confers hypersensitivity to a subset of CLE peptides. (A-B) Representative images and quantification as box plot and jitter box graphs of the root growth between TPLATE_1 and WDXM2_1 seedlings grown for 8 days in the presence or absence of low doses (10 nM) of different CLE peptides. Scale bar = 1cm. Primary root growth was quantified for a number of seedlings (16 ≤ N ≤ 28) and statistically significant differences were observed for CLE10, CLV3 and CLE40. ***, P<0.001 (selected pairs from Welch’s ANOVA post hoc pairwise comparison with the Tukey contrasts). N.S., no significant difference. The experiment was performed twice with a similar outcome. The data represented is the quantification of one experiment where at least 3 sets of seedlings were grown on separate plates.

**Figure EV2.**
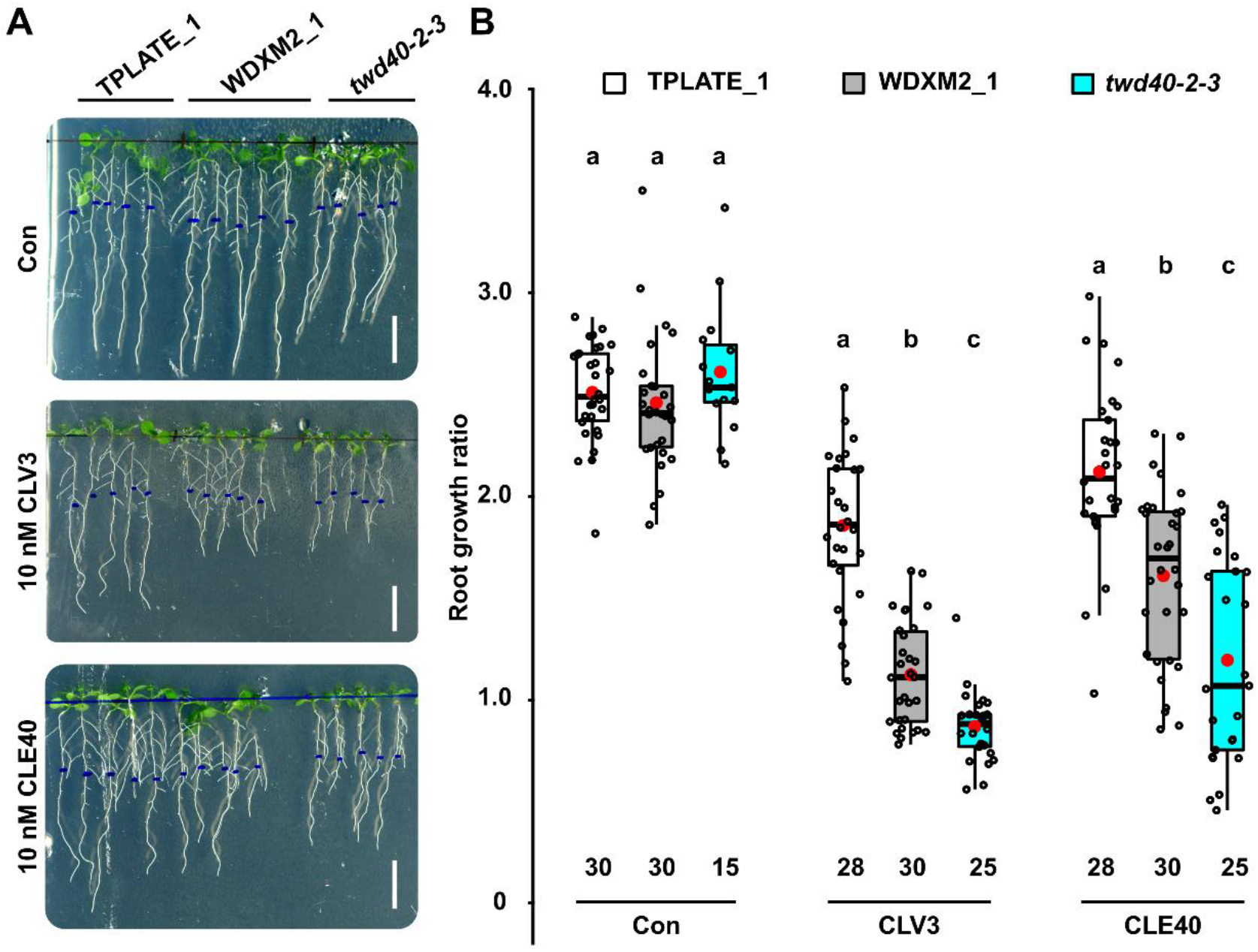
The weak TPC subunit mutant allele *twd40-2-3* confirms the observed CLV3 and CLE40 hypersensitivity in the WDXM2_1 line. (A-B) Representative images (A) and comparison (B) of the primary root growth ratios between TPLATE_1, WDXM2_1 and *twd40-2-3* seedlings transferred to plates with or without 10 nM CLV3 or CLE40. Root growth for each root after transfer was divided by the root length before transfer. Numbers of seedlings used for the quantification are indicated at the bottom of the box plot and jitter box graphs. Scale bar = 1cm. Data information in panel (B): Differences as compared to TPLATE complemented lines were evaluated by Welch’s ANOVA post hoc pairwise comparison with the Tukey contrasts. Letters (a, b and c) represent significant difference between groups (P < 0.001). The quantification represented results from at least 5 sets of seedlings grown on separate plates.

**Figure EV3.**
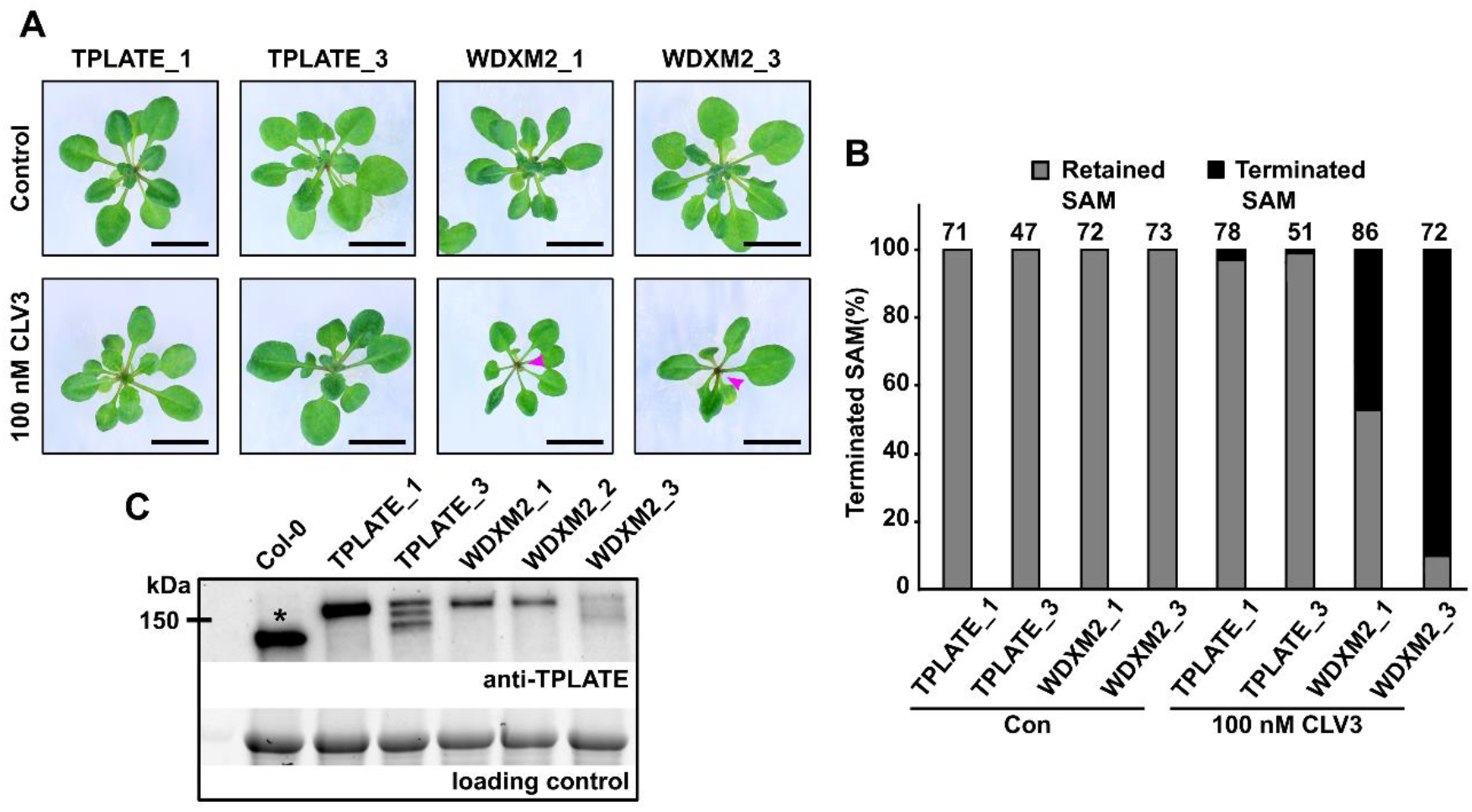
Impaired TPC-dependent endocytic capacity confers CLV3 hypersensitivity in the SAM and this hypersensitivity correlates with the expression level of the complementing construct. (A) Phenotypic comparison of 3 to 4 -week-old independent TPLATE (TPLATE_1 and TPLATE_3) and WDXM2 (WDXM2_1 and WDXM2_3) lines (GFP and mSCARLET fusions respectively) grown in the presence or absence of CLV3 peptide. Magenta arrows indicate terminated SAMs. Scale bars = 1 cm. (B) Quantification of the number of plants with a terminated meristem induced by the CLV3 peptide. The numbers of plants used for the quantification is indicated at the top of the bar chart. The experiment was repeated twice and the quantification in panel B combines both experiments. (C) Anti-TPLATE western blot detecting the presence of endogenous TPLATE in Col-0 (asterisk) as well as the full length of GFP (TPLATE _1, WDXM2_1 and WDXM2_2) or mSCARLET (TPLATE_3 and WDXM2_3) fusions of TPLATE and WDXM2 in the complemented *tplate(-/-)* homozygous mutant background that lacks endogenous TPLATE. For an unknown reason, the mSCARLET fusions give rise to several bands on the blot. The WDXM2_2 line showed a similar expression level as the WDXM2_1 and was not used further. The reduced expression in WDXM2_3 correlates with an increased hypersensitivity to the CLV3 treatment. The large subunit of RUBISCO (around 50 kDa) visualized via the stain free gel, was used as loading control.

**Figure EV4:**
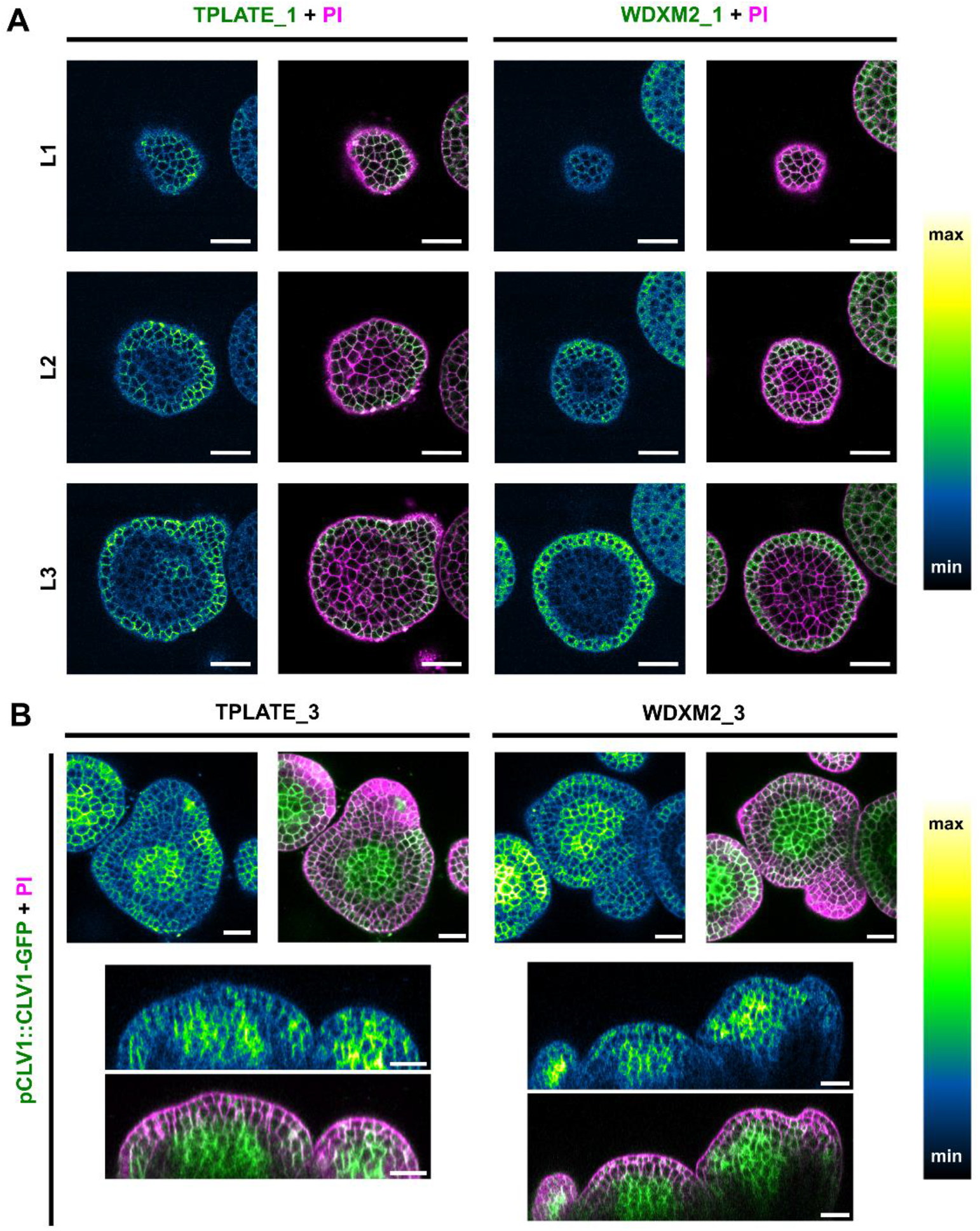
TPLATE, WDXM2 and CLV1 expression and localization in flower meristems. (A) Serial optical cross sections through the inflorescence meristem, showing localization of TPLATE and WDXM2 in layer 1 to 3 (L1-L2-L3). Left images are intensity scaled. Images on the right for each genotype represent the merged GFP and PI channels. Scale bar = 25 μm. (B) Representative confocal images showing the localization of CLV1-GFP in inflorescence meristems of TPLATE_3 and WDXM2_3 lines in optical cross sections (upper panel), and orthogonal views (lower panel). Scale bar = 20 μm.

**Figure EV5.**
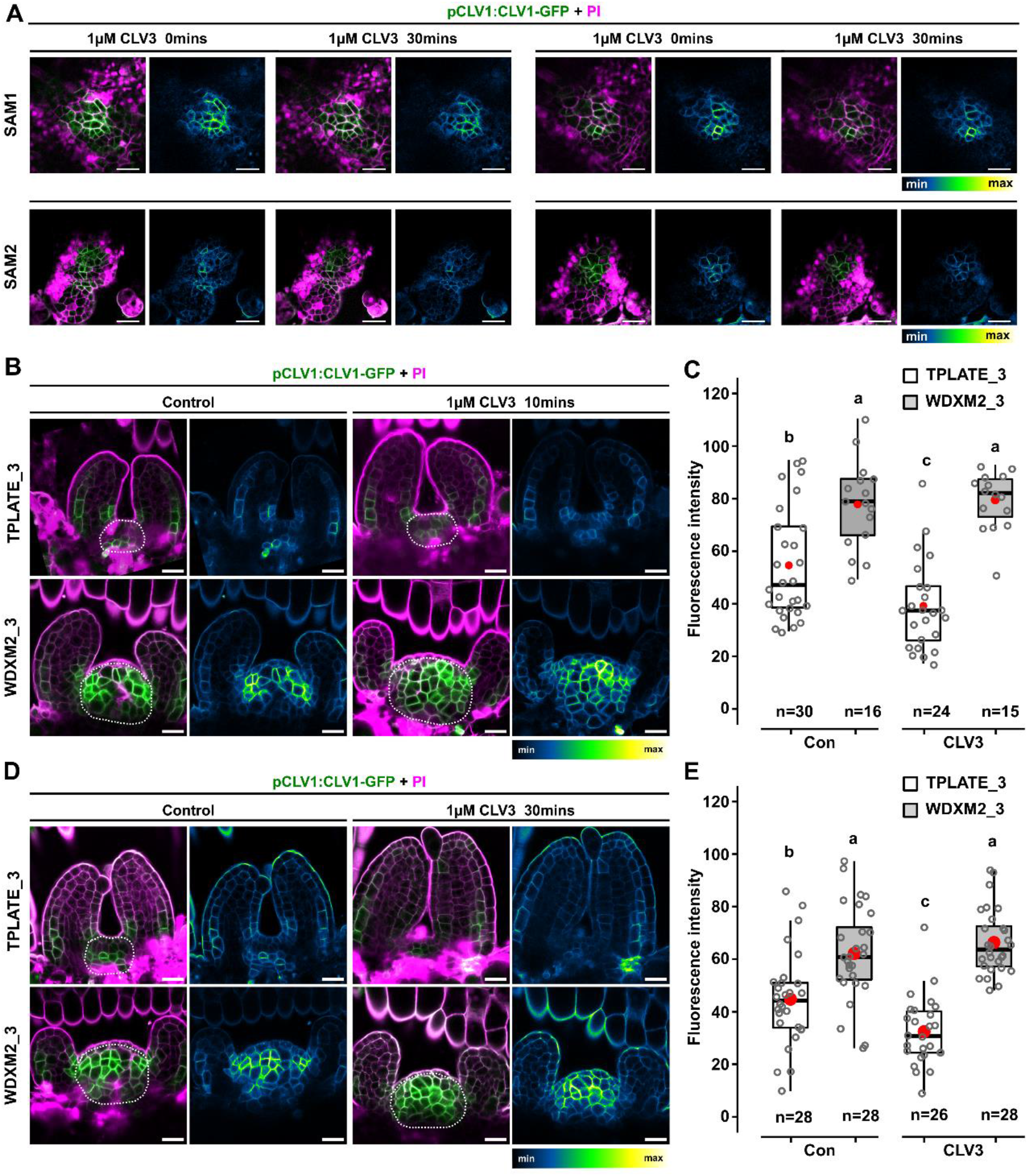
Reduced TPC-dependent endocytic capacity impairs internalization of CLV1 from PM in vegetative SAM cells upon short-term CLV3 treatment. (A) Representative images of two vegetative SAMs in the TPLATE_3 background, with different levels of CLV1 expression, imaged before and after exogenous CLV3 treatment (1 µM for 30 min). Two different focal planes of a Z-stack, visualizing different L3 cells per SAM are represented for each timepoint. For both SAMs, the CLV1 intensity at PM reduces upon CLV3 treatment. (B-E) Representative confocal images and quantification of Arabidopsis TPLATE_3 and WDXM2_3 expressing seedlings following short-term CLV3 peptide treatment (1 µM for 10 min: B-C and 1 µM for 30 min: D-E). Similar to the long-term treatment (Fig 4), PM localization of CLV1-GFP reduces upon CLV3 peptide treatment, which is not the case in WDXM2_3 vegetative meristems. The box plot and jitter box representation graph represents the average fluorescence intensity (8-bit gray values) of CLV1 over the entire SAM (indicated by a dotted line). Numbers of biological samples from two repeats are indicated at the bottom of the box plot and jitter box graphs. Differences of CLV1-GFP intensity between WDXM2_3 and TPLATE_3 lines under both conditions were evaluated by Welch’s ANOVA post hoc pairwise comparison with the Tukey contrasts. Letters (a, b and c) represent significant difference between groups (a and b, P < 0.001; b and c, P < 0.05). The quantification combines at least two independent experiments for each genotype, treatment and duration of treatment. Left panels are merged channels (GFP and PI), right panels are GFP-only channels represented via an intensity scale. Scale bar = 20 µm.

## Expanded View tables

**Table EV1:** list of used Arabidopsis lines. The table provides an overview of the lines used, their genetic background and references. H: Hygromycin; B: Basta; GPCR: Genotyping PCR; (-/-): homozygous mutant background; (+/-): heterozygous mutant background.

| Plant lines | Full name | Background | Antibiotic selection | Identification | Resources |
| --- | --- | --- | --- | --- | --- |
| *tplate* |  | (+/-) (+/+) |  | GPCR | Van Damme et al., 2006 |
| *twd40-2-3* |  | (-/-) |  | GPCR | Bashline et al., 2015 |
| **TPLATE_1** | pLAT52-TPLATE-GFP | *tplate* (-/-) | H | GPCR | Van Damme et al., 2006 |
| **TPLATE_3** | pLAT52-TPLATE-mScarlet | *tplate* (-/-) | H | GPCR | This study |
| **WDXM2_1** | pLAT52-WDXM2-GFP | *tplate* (-/-) | B | GPCR | Wang et al., 2021 |
| **WDXM2_2** | pLAT52-WDXM2-GFP | *tplate* (-/-) | B | GPCR | Wang et al., 2021 |
| **WDXM2_3** | pLAT52-WDXM2-mScarlet | *tplate* (-/-) | H | GPCR | This study |
| *clv1-101* |  | (-/-) |  |  | Kinoshita et al, 2010 |
| *clv1-8* |  | (-/-) |  |  | Dievart et al., 2003 |
| **TPLATE_1**<br>*clv1-101* | pLAT52-TPLATE-GFP | *tplate* (-/-)<br>*clv1-101* (-/-) |  | GPCR | This study |
| **WDXM2_1**<br>*clv1-101* | pLAT52-WDXM2-GFP | *tplate* (-/-)<br>*clv1-101* (-/-) |  | GPCR | This study |
| **TPLATE_1**<br>*clv1-8* | pLAT52-TPLATE-GFP | *tplate* (-/-)<br>*clv1-8* (-/-) |  | GPCR,<br>Sequence | This study |
| **WDXM2_1**<br>*clv1-8* | pLAT52-WDXM2-GFP | *tplate* (-/-)<br>*clv1-8* (-/-) |  | GPCR,<br>Sequence | This study |
| **WUS-GUS** | pWUS-GUS | *Col_0* |  |  | Su et al., 2009 |
| **TPLATE_1**<br>**pWUS-GUS** | pLAT52-TPLATE-GFP/pWUS-GUS | *tplate* (-/-) |  | GPCR, GUS staining | This study |
| **WDXM2_1**<br>**pWUS-GUS** | pLAT52-WDXM2-GFP/pWUS-GUS | *tplate* (-/-) |  | GPCR, GUS staining | This study |
| **CLV1-GFP** | pCLV1-CLV1-GFP | *Col_0* |  |  | Prof. Rüdiger Simon |
| **TPLATE_3**<br>**CLV1-GFP** | pLAT52-TPLATE-mScarlet/pCLV1-CLV1-GFP | *tplate* (-/-) |  | GPCR, Basta | This study |
| **WDXM2_3**<br>**CLV1-GFP** | pLAT52-WDXM2-mScarlet/pCLV1-CLV1-GFP | *tplate* (-/-) |  | GPCR, Basta | This study |
| **eGFP** | p35S-eGFP | *Col-0* |  |  | Obtained from Prof. E. Russinova |

**Table EV2:**
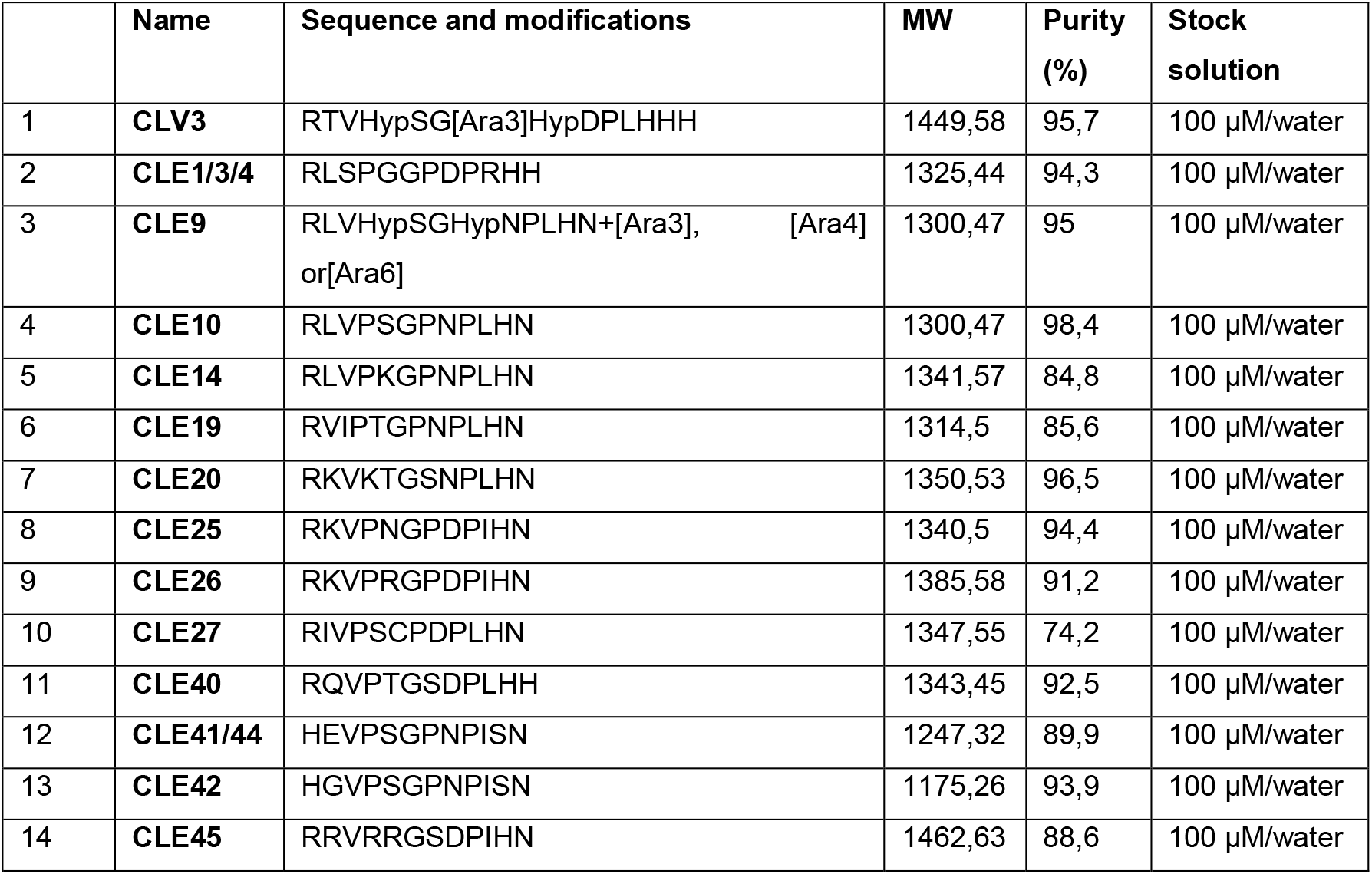
Information of CLE peptides used in this research. The table provides an overview of the sequences of the various CLE peptides used, their molecular weight (MW), purify and stock concentration. Hyp: Hydroxyproline; Ara: Arabinosylated.

## Acknowledgements

We would like to thank Elliot Meyerowitz, Marcus Heisler and Thomas Laux for constructive discussions. This work was supported by the European Research Council, Grant 682436 to D.V.D.; the Research Foundation–Flanders, Grant 1226420N to P.G.; The China Scholarship Council, grant 201508440249 to J.W.; grant 201906760018 to Q.J. and grant 201706350153 to X.X.; and by Ghent University Special Research co-funding, grant ST01511051 to J.W.

## Author contributions

J.W., Q.J., R.P. and P.G. designed and performed experiments. G.D. provided unpublished materials. C.G-A. and E. B. designed and performed floral SAM imaging. E.B performed vegetative SAM Z-stack vertical imaging. X.X. and P.G. performed co-IP. M.V. performed Y2H. E.M. performed confocal imaging. R.P., I.D.S., T.V., R.S., M. K. N. and D.V.D. designed and supervised research. J.W., R.P., and D.V.D wrote the initial draft of the manuscript. All authors were involved in discussing the data and in finalizing the conclusions and text of the manuscript.

## Data availability

This study includes no data deposited in external repositories. Data used for quantifications as well as full Western blots can be found in the source data file. All material will be made available upon reasonable request to the corresponding author.

